# The role of homology and orthology in the phylogenomic analysis of metazoan gene content

**DOI:** 10.1101/341115

**Authors:** Walker Pett, Marcin Adamski, Maja Adamska, Warren R. Francis, Michael Eitel, Davide Pisani, Gert Wörheide

## Abstract

Resolving animal (Metazoa) relationships is crucial to our understanding of, for example, the origin of their key traits such as muscles, guts and nerves. However, a broadly accepted metazoan consensus phylogeny has yet to emerge. In part this is because the genomes of deeply-diverging and fast-evolving lineages may undergo significant gene turnover, reducing the number of orthologs shared with related phyla. This can limit the usefulness of traditional phylogenetic methods that rely on alignments of orthologous sequences. Phylogenetic analysis of gene content has the potential to circumvent this orthology requirement, with binary presence/absence of homologous gene families representing a source of phylogenetically informative characters. Applying binary substitution models to the gene content of 26 complete animal genomes, we demonstrate that patterns of gene conservation differ markedly depending on whether gene families are defined by orthology or homology, i.e., whether paralogs are excluded or included. We conclude that the placement of deeply-diverging lineages, like ctenophores, may exceed the limit of resolution afforded by methods based on comparisons of orthologous protein supermatrices, and novel approaches are required to fully capture the evolutionary signal from genes within genomes.

## Introduction

Resolving the phylogenetic relationships among animal lineages is crucial for understanding, for example, the evolutionary origin of key animal traits, such as a true gut, muscles and a nervous system. With advancements in sequencing technology, and the associated increase in the availability of genomic data from these groups, especially the phylogeny of early animals has attracted renewed attention and has been the focus of many recent phylogenomics studies (summarized in Dunn et al. 2014; King and Rokas 2017). In particular, the phylogenetic position of Ctenophora has been disputed by numerous authors in recent years (Philippe et al. 2009; Pick et al. 2010; Nosenko et al. 2013; Pisani et al. 2015; Whelan et al. 2015; Feuda et al. 2017; Simion et al. 2017; Whelan et al. 2017). Previous studies focusing on deep metazoan phylogeny have relied mostly on the analysis of amino acid sequences from a large number of proteins, either concatenated into a single data matrix (“supermatrix”) (Philippe et al. 2009; Ryan et al. 2013; Moroz et al. 2014; Whelan et al. 2015; Shen et al. 2017; Simion et al. 2017), or separately as individual gene family alignments (Arcila et al. 2017). These methods generally rely on the identification of sequences that are related by orthology (orthologs), whose common ancestor diverged as a result of speciation, rather than a duplication event (Fitch 2000). Genes arising from duplication events (paralogs) complicate the inference of a species tree based on concatenated gene alignments. This is because the gene tree describing the relationships among paralogs may differ markedly from the species tree (Martin and Burg 2002; Struck 2013). Including paralogous sequences in a phylogenetic analysis of a “supermatrix” can therefore lead to artifacts in reconstructing a species phylogeny, and various methods aimed at removing paralogs from amino acid datasets have been developed (Li et al. 2003; Fulton et al. 2006; van der Heijden et al. 2007; Gabaldón 2008; Pereira et al. 2014; Struck 2014). One of the most popular of ortholog selection approaches is OrthoMCL (Li et al. 2003), which uses a reciprocal best hits (RBH) Blast algorithm (all vs all) for ortholog identification, combined with the Markov clustering algorithm (MCL) (Van Dongen 2001; Enright et al. 2002) to find clusters of orthologous genes.

Besides amino acid sequences, the presence or absence of genes in different genomes has been suggested as an alternative source of phylogenetically informative genomic characters (Fitz-Gibbon and House 1999; Lake and Rivera 2004; Ryan et al. 2013; Pisani et al. 2015). However the issue of identifying genes related by paralogy and orthology is still an important consideration for the analysis of gene content, since the presence of a gene family in a species does not necessarily imply that all orthologous subfamilies are also present. Thus, we may consider independently both homolog (i.e. both paralogs and orthologs) and ortholog content in phylogenetic reconstruction.

Here we present a phylogenetic analyses of animals using gene content inferred from complete genomes. We analyse datasets scoring both orthology groups and protein families. We show that relationships inferred from both type of data are highly congruent, only differing in the relative relationships of the Ctenophora, that emerge as the sister group of all animals but the sponges (Porifera-sister hypothesis) when presence/absence of orthology groups are analysed, or as the sister of the Cnidaria (Coelenterata Hypothesis) when presence/absence of protein families is analysed. Furthermore, our results provide insights into the behavior of different gene content datasets that may help direct future investigations using this type of data.

## Materials and Methods

### Proteome data acquisition

We began by reconstructing the gene content dataset of Ryan et al. (2013), which included 23 complete genomes, with 21 from across metazoa, including one ctenophore (*Mnemiopsis leidyi*) and one sponge (*Amphimedon queenslandica*), as well as 2 complete genomes from unicellular relatives of animals (*Capsaspora owczarzaki* and *Monosiga brevicollis*). We obtained predicted complete proteomes for each of these genomes either from the Ensembl Metazoa database (Cunningham et al. 2015), or from the Origins of Multicellularity Database hosted by the Broad Institute (Ruiz-Trillo et al. 2007). In addition to this dataset of 23 species (Meta23), we constructed an expanded dataset which included 13 additional proteome predictions based on complete genomic data (Meta36), including the ctenophore *Pleurobrachia bachei* from the Neurobase genome database (Moroz et al. 2014), the homoscleromorph sponge *Oscarella carmela* from the Compagen database (Hemmrich and Bosch 2008), as well as 4 fungal species from the Ensembl database (Cunningham et al. 2015) and 2 species of fungi and two unicellular relatives of animals from the Origins of Multicellularity Database (Ruiz-Trillo et al. 2007).

### Proteome prediction

In addition we included new proteome assemblies predicted from complete genomic and transcriptomic data for the calcareous sponge *Sycon ciliatum*, as well as newly described placozoan species from a new genus (Eitel et al. 2017). Details on Placozoa sp. H13 (Eitel et al. 2013) genome sequencing and annotation can be found elsewhere (Eitel et al. 2017). The *Sycon ciliatum* transcriptome was assembled de novo from a comprehensive set of Illumina RNA-Seq libraries using Trinity pipeline (Grabherr et al. 2011). The libraries (PE, poly-A) were generated from *S. ciliatum* larvae, various developmental stages and regenerating adult specimens, ENA submissions ERA295577 and ERA295580 (Fortunato et al. 2014; Leininger et al. 2014). Protein-coding sequences (CDS’es) were detected with Transdecoder (Haas et al. 2013) using Pfam database (Finn et al. 2016) with minimum length cut-off of 300 nucleotides. To remove potential microbial contaminations the amino-acid translations of the CDS’es were BLASTP searched against NCBI NR protein database. All hits with better score to archaea, bacteria or viruses than to eukaryotes were removed. Remaining CDS’es were clustered with CDHIT (Li et al. 2001) with parameters -G 1 -c 0.75 -aL 0.01 -aS 0.5.

### Gene family prediction

For each dataset, we computed two clusterings based on either homology (as defined by a Blast e-value threshold) or orthology (as defined by OrthoMCL). This resulted in four final clusterings (Meta23-Homo, Meta36-Homo, Meta23-Ortho and Meta36-Ortho). To build the clusterings, we performed an all vs. all BlastP query with all protein sequences, with a minimum e-value cutoff of 1e-5, keeping all hits and high-scoring segment pairs (HSPs) for each query sequence. These Blast results were used directly as input to OrthoMCL (Li et al. 2003) for the prediction of orthologous gene clusters. For the prediction of homologous gene clusters, we directly transformed the BlastP output into an edge-weighted similarity graph, using a weighting algorithm identical to that used by OrthoMCL, with the only difference being that the reciprocal best hits (RBH) and normalization steps were skipped, the functions of which are to discriminate orthologous and paralogous pairwise relationships. We implemented this modified OrthoMCL algorithm in C++, the source code for which has been deposited on GitHub (http://www.github.com/willpett/homomcl). Exactly as in OrthoMCL, we computed edge weights as the average -log_10_ e-values of each Blast query-target sequence pair, where e-values equal to 0.0 were set to the minimum e-value exponent observed across all pairs. Also as in OrthoMCL, we computed the “percent match length” (PML) for each pair as the fraction of residues in the shorter sequence that participate in the alignment, and used the default PML cutoff of 50% for each edge (see the OrthoMCL algorithm document for more details). Using these edge-weighted similarity graphs, clusterings were computed using the Markov clustering algorithm (MCL) (Van Dongen 2001) with an inflation parameter of 1.5, which is the default suggested by OrthoMCL. The structures of resulting clusterings were compared using the clm info and clm dist commands in the mcl package.

### Phylogenetic analysis

For each of the four clusterings, a presence/absence matrix was constructed where each species was coded as present or absent in a cluster depending on whether at least one protein sequence from that species was contained in that cluster. Phylogenetic trees were reconstructed from the resulting binary data matrices in RevBayes (Höhna et al. 2015) using the reversible binary substitution model of (Felsenstein 1992; Ronquist et al. 2012). We also used an irreversible Dollo substitution model (Nicholls and Gray 2006; Alekseyenko et al. 2008), in which each gene family may be gained only once, and thereafter follows a pure-loss process. We computed the marginal likelihood for each model using stepping-stone sampling as implemented in RevBayes. In all cases, we used 4 discrete categories for gamma-distributed rates across sites, and we used a hierarchical exponential prior on the branch lengths. RevBayes scripts for these analyses have been deposited on GitHub (http://www.github.com/willpett/metazoa-gene-content). Convergence statistics were computed using the programs bpcomp and tracecomp in the PhyloBayes package (Lartillot et al. 2013), and are reported in Supplementary Table S1.

### Correcting for unobserved losses

Importantly, many clusters were observed in only a single species, which we define as “singleton” gene families. The singleton gene families identified in our analysis represent only a subset of all singletons, because some cannot be observed. For example, genes without significant sequence similarity to other sequences may not have been identified in the original proteome prediction. Furthermore, many singleton gene families represented by a single sequence, which we define as “orphans”, may not have had any significant BlastP hits in the all vs. all query. Therefore, in order to avoid introducing an ascertainment bias by including only a subset of singletons (Pisani et al. 2015; Tarver et al. 2018), we removed all singletons from each data matrix prior to phylogenetic analysis, and applied a correction for the removal of singletons in RevBayes (coding=nosingletonpresence). To further evaluate the impact of including/excluding singletons, we approximated the total number of singleton families as the observed number singleton clusters plus the number of orphans. In an additional analysis, we included these orphans along with the previously-excluded observed singletons without applying the “nosingletonpresence” ascertainment bias correction. Finally, in all analyses we also applied a correction for the fact that genes absent in all species cannot be observed (coding=noabsencesites).

### Posterior Predictive Simulations

We evaluated the impact of including singletons in our phylogenetic analysis by simulating *n_0_*, the number of gene families present in 0 species (lost in all species), and *n_1_*, the number of gene families present in 1 species (singletons), from their respective posterior predictive distributions, either conditioned on the inclusion or exclusion of singletons. Specifically, for each sample *θ* from the posterior, and given the observed number of gene families present in more than one species *N*, we simulated *n_0_* and *n_1_* from the predictive distribution

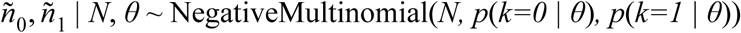

where *p*(*k|θ*) is the binary substitution model likelihood of observing a gene family present in *k* species. The observed values for *n_1_* and *N* for each analysis are listed in Table 1. Note that *n_0_* cannot be observed. Simulations were implemented in biphy, a software package for phylogenetic analysis of binary character data (http://www.github.com/willpett/biphy).

**Table 1.** Clustering statistics for homologous and orthologous gene family predictions in two datasets

| **Dataset** | **Method** | **Edges** | **Orphans** | **Clusters** | **Singletons** | **$N$** | **$n_1$** | **% Sub. Diff.** |
| --- | --- | --- | --- | --- | --- | --- | --- | --- |
| meta23 | Homologs | 50096494 | 74021 | 22679 | 10596 | 12083 | 84617 | NA |
| meta36 | Homologs | 73588623 | 116566 | 35244 | 17513 | 17731 | 134079 | NA |
| meta23 | Orthologs | 4468075 | 118103 | 39240 | 16955 | 22285 | 135058 | 0.029 |
| meta36 | Orthologs | 6445128 | 175913 | 57056 | 25458 | 31598 | 201371 | 0.033 |

## Results and Discussion

### Ortholog content vs homolog content in phylogenetic reconstruction

Using a reciprocal best hits (RBH) algorithm, OrthoMCL aims to identify pairs of proteins related by either orthology, co-orthology, or in-paralogy. This is accomplished using a similarity graph based on BlastP e-values, from which edges that OrthoMCL identifies as representing pairwise paralogous relationships are removed. Thus, since most pairs of homologous sequences are related by paralogy, the vast majority of edges may be removed from the full similarity graph using this algorithm. Indeed, in our analysis, the OrthoMCL RBH algorithm resulted in the removal of 91% of edges from both graphs (Table 1). Moreover, removing these edges had a major impact on the granularity of the resulting MCL clusterings, with nearly twice as many clusters identified after paralogous edges had been removed (Table 1). In addition, these clusterings differed by only about 3% from a sub-clustering of the one obtained on the full graph (Table 1). This is indeed the goal of OrthoMCL: to identify sub-clusters of orthologous sequences, which by definition outnumber clusters of sequence that are merely homologous.

However, by subdividing clusters of homologous sequences into orthologous groups, some phylogenetic signal regarding the presence or absence of homologous gene families may be removed from the final presence/absence matrix. Indeed, our estimates for the expected number of gain/loss events per gene family across metazoa (the tree length) were smaller in the analysis of homologs (posterior mean = 0.065) compared to orthologs (posterior mean = 0.172), confirming the expectation that homolog content is more strongly conserved than ortholog content in animals The relative rate of gene family loss was also several-fold smaller in the analysis of homolog content (posterior mean = 0.0012) compared to ortholog content (posterior mean = 0.0026).

We began with a phylogenetic analysis of the reconstructed dataset of Ryan et al. (2013) using OrthoMCL (Meta23-Ortho), and then comparing it to the analogous dataset of homologous gene family content based on BlastP similarity only (Meta23-Homo). Phylogenies constructed based on ortholog and homolog content differed only in the position of Ctenophora. As in Pisani et al. (2015), when the data matrix represented the presence or absence of orthogroups identified by OrthoMCL, strong support was obtained for the a tree where the sponges represented the sister group of all the other animals. In this tree, Ctenophora was found to represent the sister group of all the animals but the sponges (Porifera-sister hypothesis; Fig S1). When the data represented presence or absence of homologous clusters (paralogous relationships had not been removed by OrthoMCL), sponges were still found to represent the sister of all the other animals. However, this analysis recovered Ctenophora as the sister group of Cnidaria plus Bilateria, albeit with relatively weak support (Fig S2).

We then expanded the taxon sampling of our dataset to incorporate thirteen additional genomes, including 5 non-bilaterian animals representing each major non-bilaterian phylum, as well as 8 non-metazoan outgroup species, ranging in evolutionary distance from fungi to choanoflagellates. Similarly to Meta-23-Ortho, this dataset (Meta36-Ortho), gave strong support for the sponges as the sister group of all the other animals and Ctenophora as the sister to all non-Poriferan animals (Porifera-sister hypothesis; Fig. S3). Differently, analysis of homolog content (Meta36-Homo) gave strong support to Ctenophora as the sister group of Cnidaria (i.e. the Coelenterata hypothesis; Fig 1). This result was not sensitive to the choice of outgroup, and was strongly supported even when 6 species of fungi were included.

**Figure 1.**
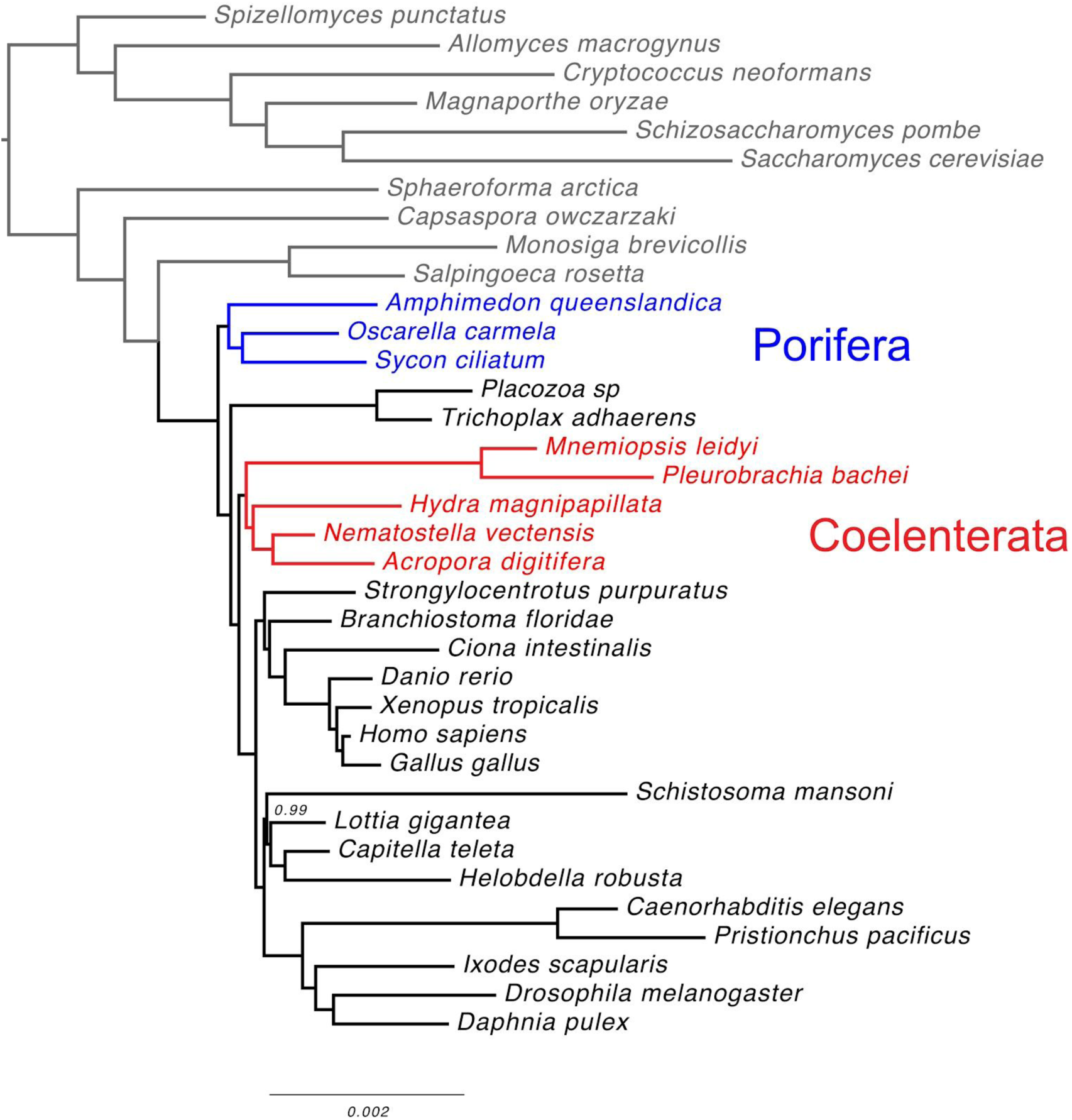
Phylogenetic tree reconstructed from Bayesian analysis of homologous gene family content in 36 species of opisthokonts, including 26 animals (Meta36-Homo). Singletons were excluded, and an ascertainment bias correction was applied for the absence of singletons and gene families lost in all species. Numbers of nodes indicate posterior probabilities indistinguishable from 1.0. Convergence statistics are listed in Supplementary Table S1.

The placement of Ctenophora outside of Eumetazoa is found only by the analysis of ortholog content, which is consistent with other researchers’ previous findings that many eumetazoan-specific orthologs are absent in ctenophores (Ryan et al. 2010; Ryan et al. 2013; Moroz et al. 2014). Furthermore, the support for Coelenterata based on homolog content suggests that while many eumetazoan and coelenterate orthologs may have been lost in ctenophores, related paralogs specific to Ctenophora may have been retained in the same gene families. Alternatively, these putative paralogs may represent the missing eumetazoan orthologs which have evolved to such an extent that OrthoMCL cannot reliably identify them as orthologs anymore. This apparent absence of Eumetazoa-specific orthologs in Ctenophora may help to explain the great difficulty that previous phylogenomic studies based on concatenated amino acid sequences of orthologous proteins have had in resolving the position of ctenophores. If eumetazoan gene families are indeed represented in ctenophores mostly by sequences that are, or appear to be, paralogous to other eumetazoans, then these gene families will be systematically removed during the construction of of orthologous amino acid sequence alignments, resulting in relatively little signal for a eumetazoan affinity of Ctenophora.

### Singleton gene families

The issue of whether to include singleton gene families in the analysis of gene content is complicated by the fact that the number of singletons is directly correlated with the stringency of our definition of homology. By increasing the e-value cutoff during the Blast query step, an arbitrary number of protein sequences can be excluded as lacking significant similarity to any other sequences. Including these orphans can therefore lead to an overestimate of the actual number of singleton gene families. On the other hand, excluding orphans will lead to an underestimate of the number of singletons. To demonstrate this effect, we estimated the posterior predictive distribution of the number of singleton gene families predicted by the binary substitution model after either including singletons and orphans, or excluding both (Figs 2-3, S7-8). In both cases, the predicted number of singleton families was much smaller than the observed combined number of singletons and orphans, suggesting that the number of orphans is indeed an overestimate of the actual number of unobserved singleton gene families. In fact, including orphans appears to introduce an upward bias in the estimation of the gene family loss rate, such that the predicted number of singletons actually decreases when observed singletons are included in the analysis (Fig 3), inflating the predicted number of gene families lost in all species (Fig 2). Faced with this inherent bias in estimating the number of singletons, it is perhaps unsurprising that our phylogenetic analysis including singletons recovered an unconventional tree with Placozoa as the sister group to all other animals (Fig S4-5). We conclude that simply excluding all singletons and applying an appropriate correction provides less biased estimates of gene gain and loss rates, and consequently the tree topology.

**Figure 2.**
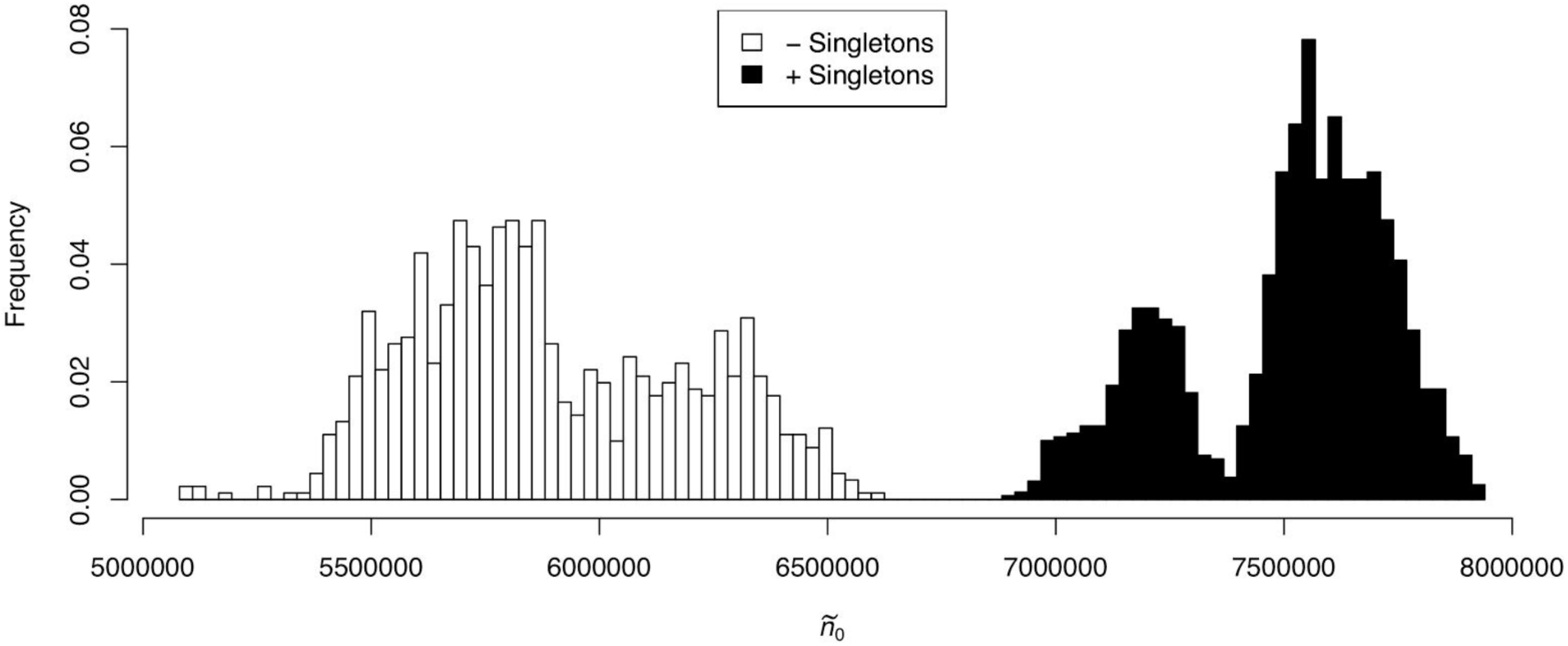
Posterior predictive distribution of the number of homologous gene families lost in all species (ñ_0_), inferred from Meta36-Homo using a reversible binary substitution model, either including singletons (black) or excluding them (white).

**Figure 3.**
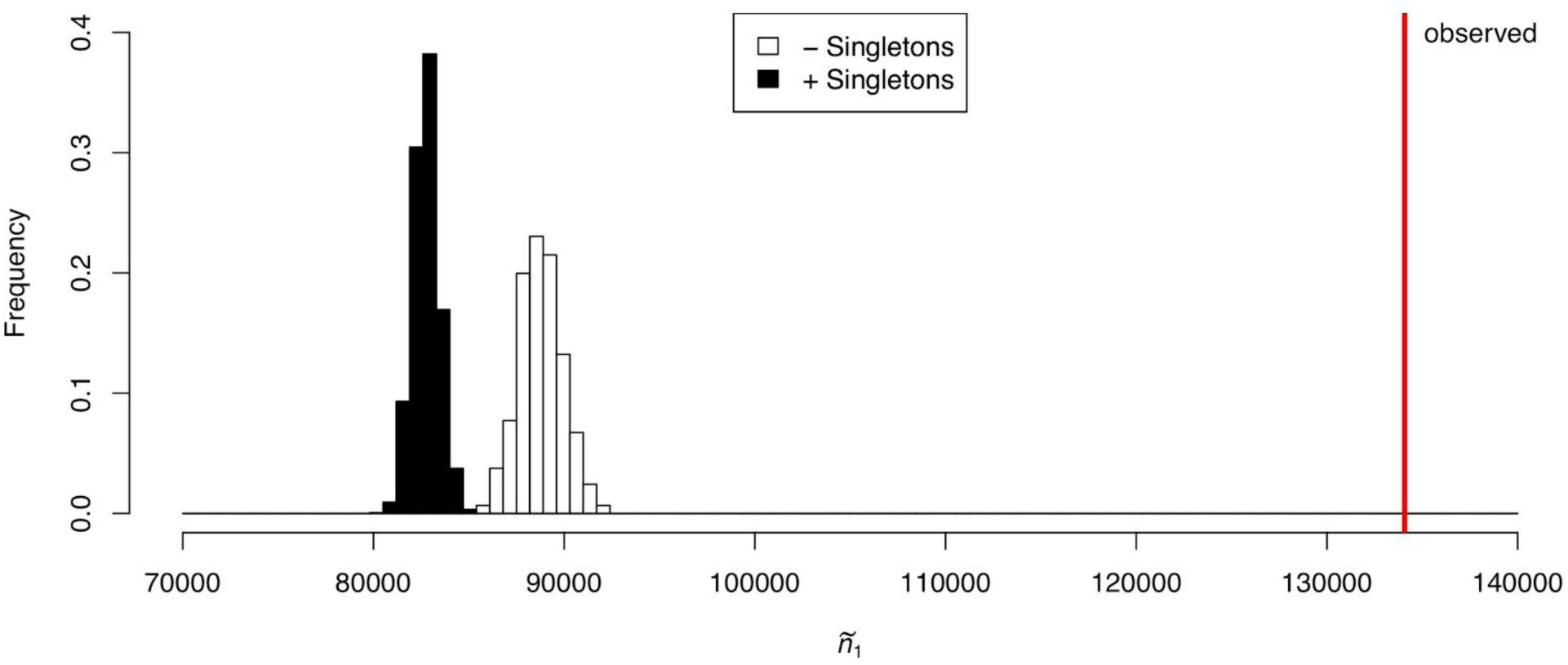
Posterior predictive distribution of the number of homologous gene families present in only a single species (ñ_1_), inferred from Meta36-Homo using a reversible binary substitution model, either including singletons (black) or excluding them (white).

### Weak support for irreversible gene content evolution in animals

We found that a reversible binary substitution model (Felsenstein 1992), in which a gene family may be gained more than once on the tree, provided a much better fit to the Meta36-Homo dataset than an explicitly irreversible Dollo-like model (Nicholls and Gray 2006; Alekseyenko et al. 2008), in which each gene family may be gained only once (Table 2). Previous authors have interpreted similar results as evidence for horizontal gene transfer (HGT) in prokaryotes (Zamani-Dahaj et al. 2016). However, only a few cases of HGT among animals have been reported (Jackson et al. 2011; Boto 2014). Therefore, we find horizontal gene transfer an unlikely explanation for the stronger fit of a reversible model in our analysis of animal genomes. Furthermore, both the reversible and Dollo models recovered identical tree topologies (Fig. 1 and S6 respectively). This suggests that the source of the reversible model’s improved fit is not underlying phylogenetic signal, but may instead represent noise related to errors in the prediction of gene family clusters, or in the prediction and assembly of protein sequence databases.

**Table 2.** Marginal likelihoods estimated for the meta36-Homo dataset

| **Model** | **Marginal Likelihood** |
| --- | --- |
| Reversible | -174066 |
| Dollo | -179849 |

## Conclusion

The construction of any phylogenomic dataset entails considerable difficulties in the identification of a sufficient number of homologous characters to infer a well-resolved species tree. Methods that utilize a single concatenated alignment of multiple loci to infer a species tree are further limited by the requirement for orthologous sequences, as this is the only way to ensure that a single tree describes their evolutionary history. Using gene content, or the presence or absence of gene families, as character data can circumvent this orthology requirement and may therefore allow inferences at deeper time-scales. Our phylogenetic analysis of homologous, but not orthologous gene family content data from 26 metazoan species and 10 non-metazoans strongly supports the classical view of animal phylogeny, with sponges as the sister group to all other animals. We conclude that the placement of some deeply-diverging lineages, like ctenophores, may exceed the limit of resolution afforded by traditional methods that use concatenated alignments of individual genes, and novel approaches are required to fully capture the evolutionary signal from genes within genomes. Finally, our analysis shows that accounting for ascertainment bias in gene family size (i.e. widely distributed gene families are sampled at a greater rate than those with small taxonomic range) is of general importance in the future development of probabilistic models of gene family evolution.

## Acknowledgements

This work was supported by funding by a NERC BETR (NE/P013643/1) Grant to DP, as well as from the LMU Munich’s Institutional Strategy LMU excellent within the framework of the German Excellence Initiative, and the German Research Foundation (DFG) grant Wo896/19-1, and from the European Union’s Horizon 2020 research and innovation programme under the Marie Skłodowska-Curie grant agreement No 764840 (ITN IGNITE) to GW.

---

**Figure S1.**
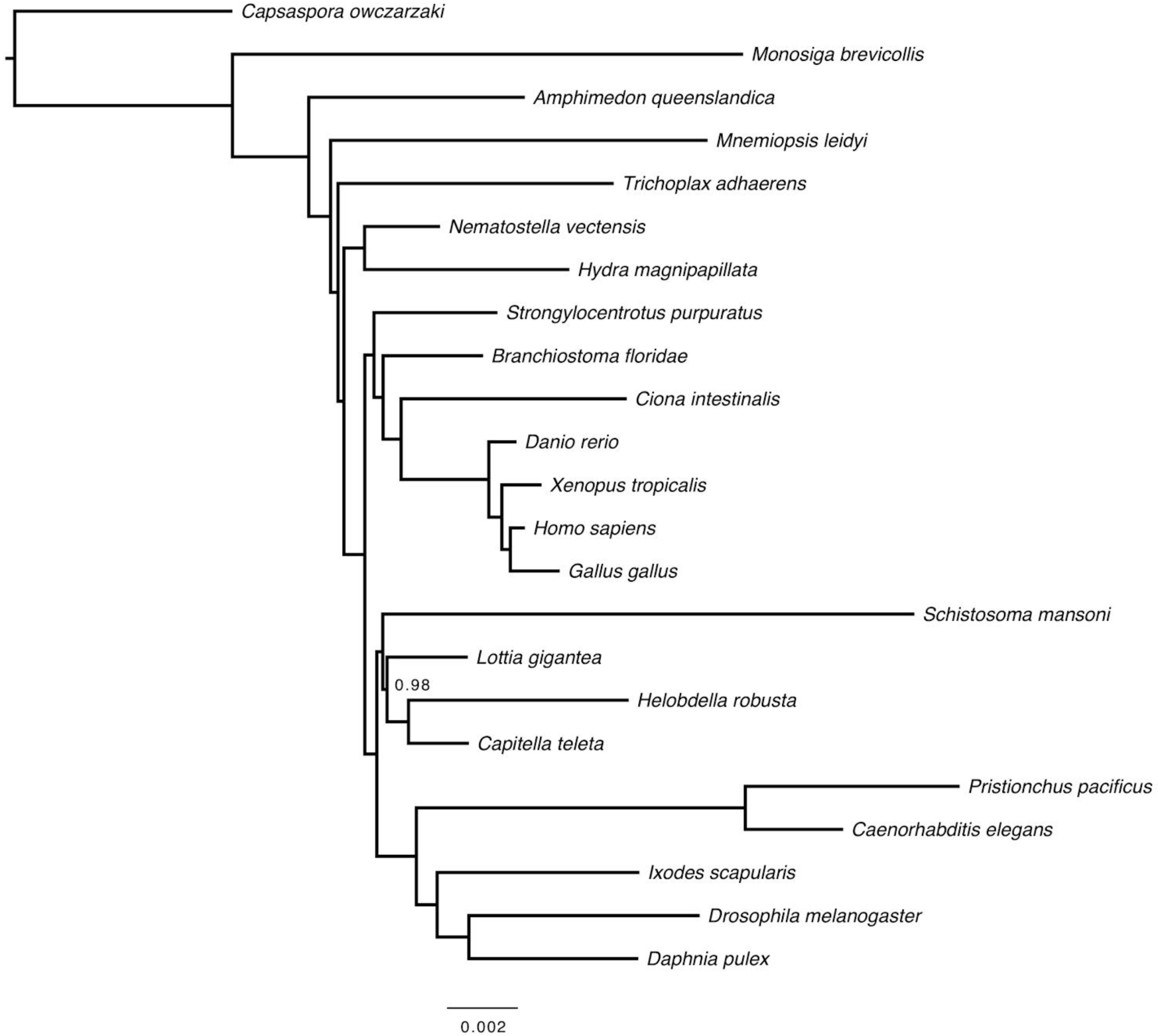
Phylogenetic tree reconstructed from Bayesian analysis of orthologous gene family content in 23 species of opisthokonts, including 21 animals (Meta23-Ortho) using a reversible binary substitution model. Singletons were excluded, and an ascertainment bias correction was applied for the absence of singletons and orthologs lost in all species. Numbers of nodes indicate posterior probabilities indistinguishable from 1.0. Convergence statistics are listed in Supplementary Table S1.

**Figure S2.**
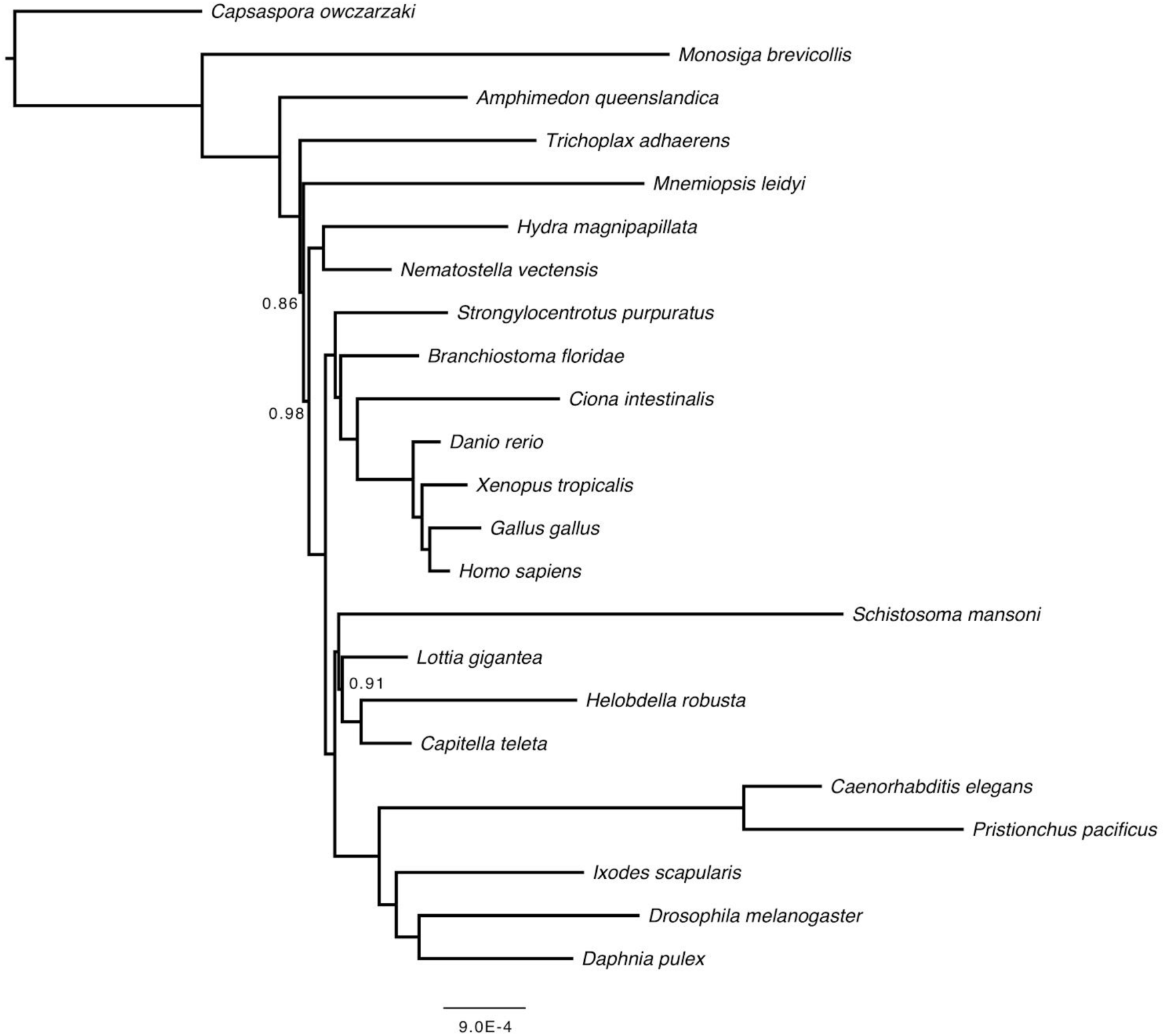
Phylogenetic tree reconstructed from Bayesian analysis of homologous gene family content in 23 species of opisthokonts, including 21 animals (Meta23-Homo) using a reversible binary substitution model. Singletons were excluded, and an ascertainment bias correction was applied for the absence of singletons and gene families lost in all species. Numbers of nodes indicate posterior probabilities indistinguishable from 1.0. Convergence statistics are listed in Supplementary Table S1.

**Figure S3.**
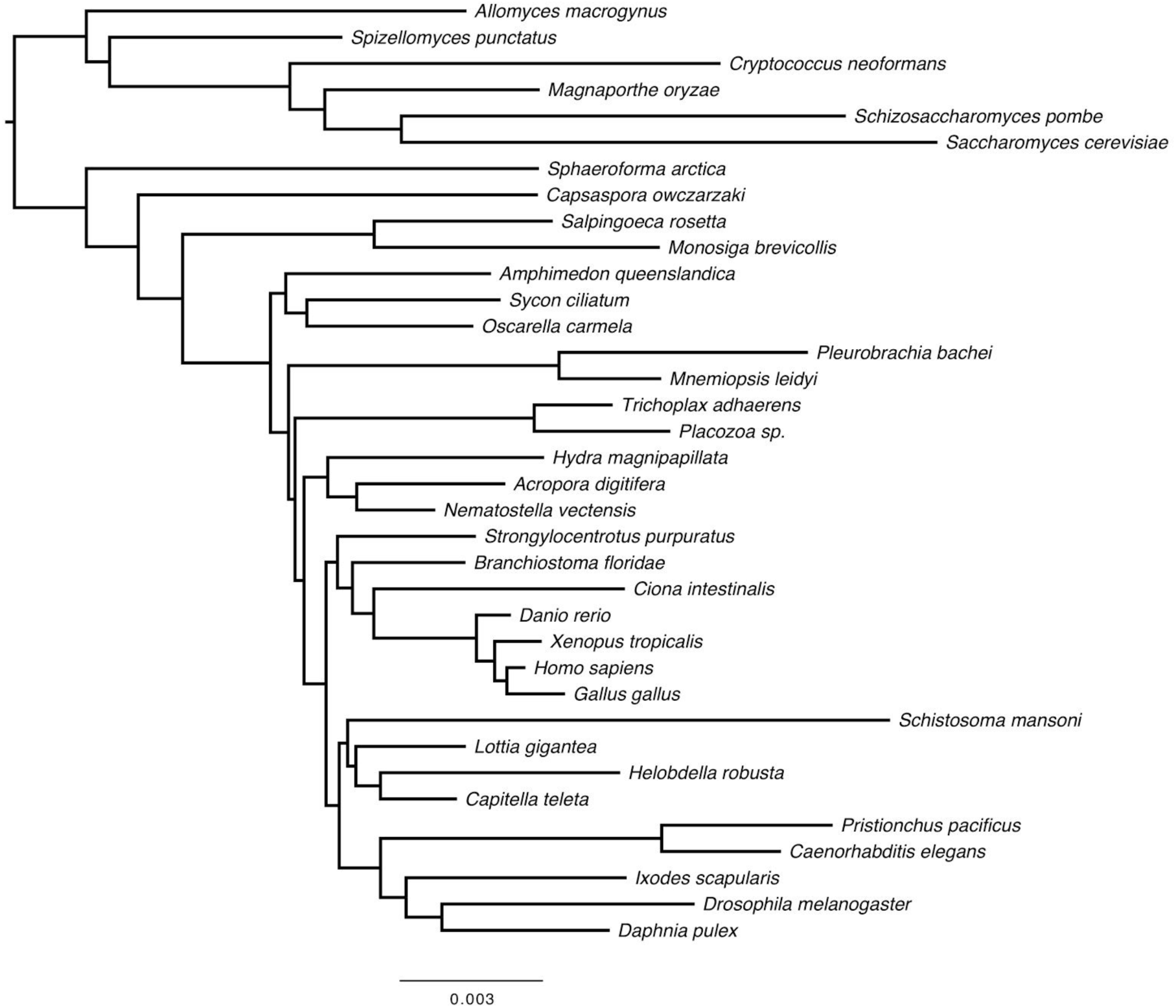
Phylogenetic tree reconstructed from Bayesian analysis of orthologous gene family content in 36 species of opisthokonts, including 26 animals (Meta36-Ortho) using a reversible binary substitution model. Singletons were excluded and an ascertainment bias correction was applied for the absence of singletons and orthologs lost in all species. Numbers of nodes indicate posterior probabilities indistinguishable from 1.0. Convergence statistics are listed in Supplementary Table S1.

**Figure S4.**
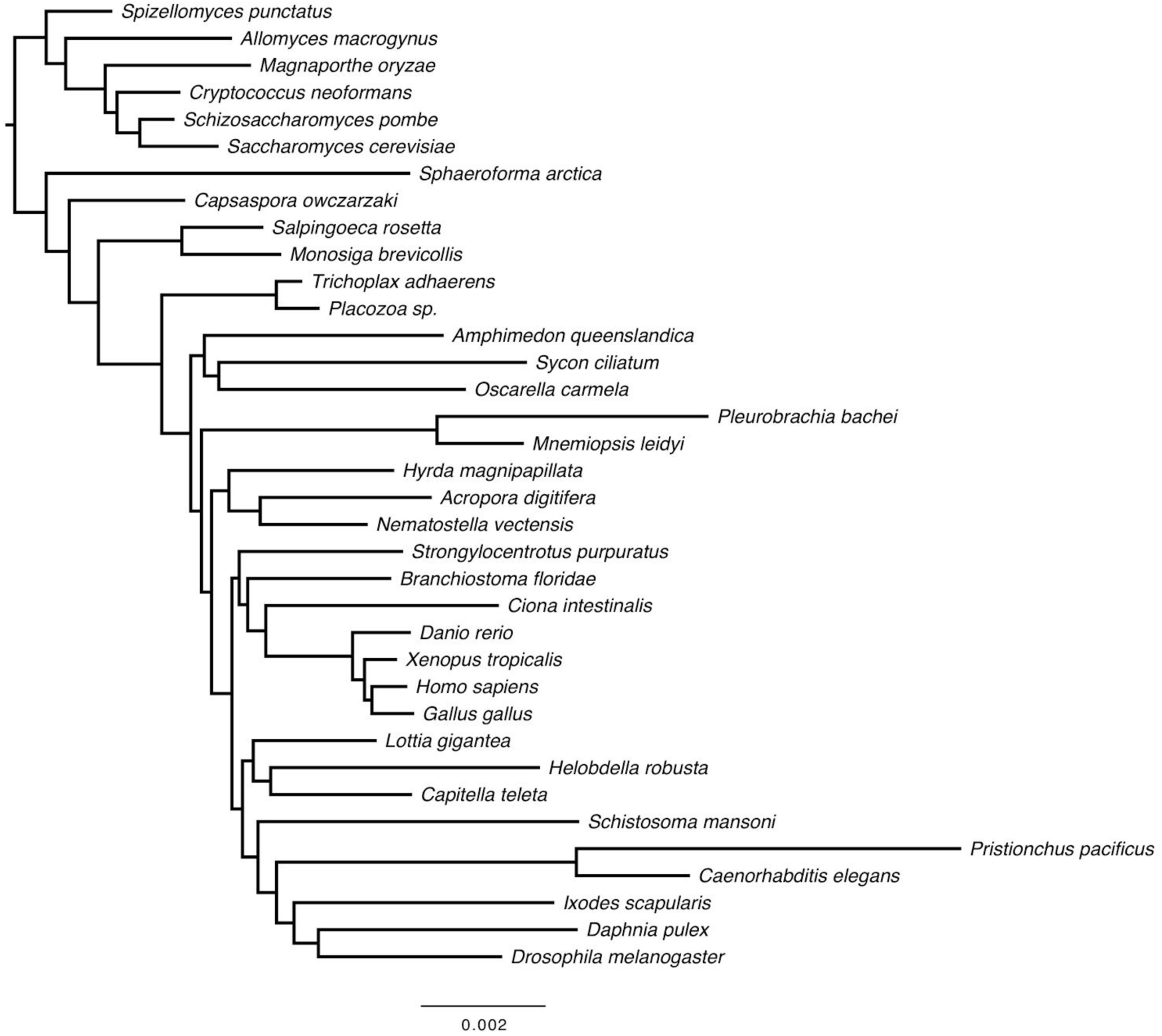
Phylogenetic tree reconstructed from Bayesian analysis of orthologous gene family content in 36 species of opisthokonts, including 26 animals (Meta36-Ortho) using a reversible binary substitution model. Singletons and orphans were included, and an ascertainment bias correction was applied for the absence gene families lost in all species. Numbers of nodes indicate posterior probabilities indistinguishable from 1.0. Convergence statistics are listed in Supplementary Table S1.

**Figure S5.**
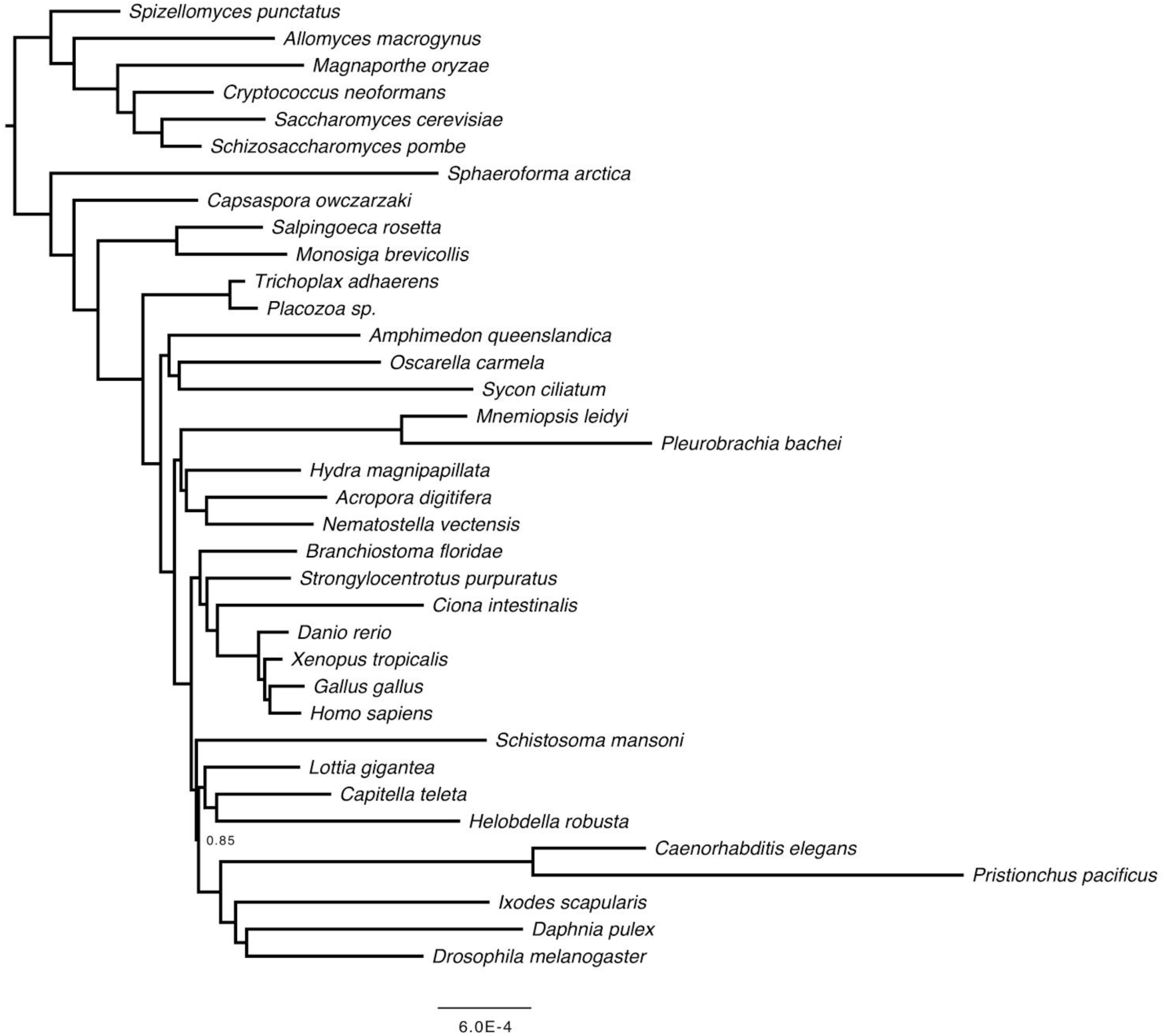
Phylogenetic tree reconstructed from Bayesian analysis of orthologous gene family content in 36 species of opisthokonts, including 26 animals (Meta36-Homo) using a reversible binary substitution model. Singletons and orphans were included, and an ascertainment bias correction was applied for the absence gene families lost in all species. Numbers of nodes indicate posterior probabilities indistinguishable from 1.0. Convergence statistics are listed in Supplementary Table S1.

**Figure S6.**
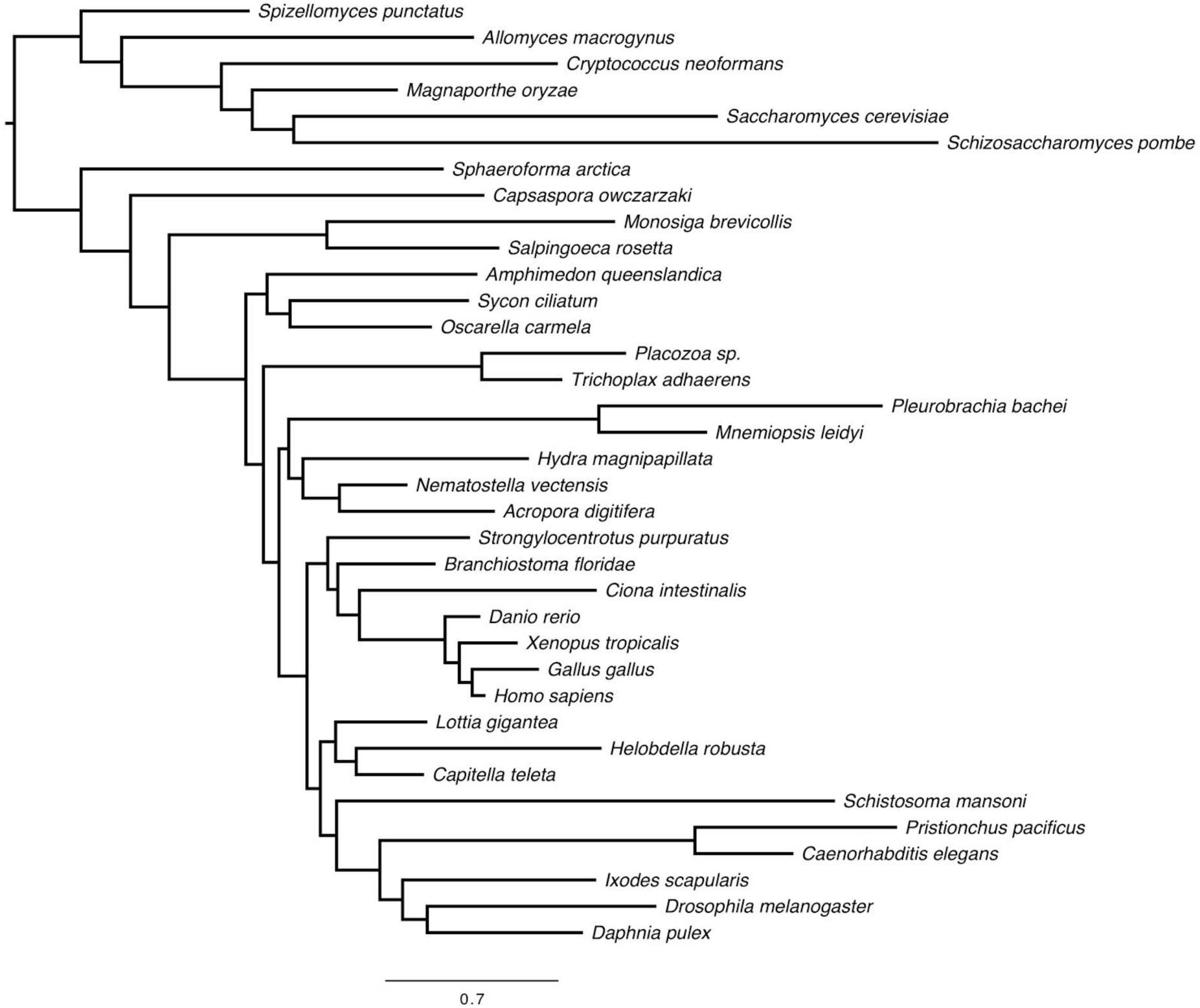
Phylogenetic tree reconstructed from Bayesian analysis of homologous gene family content in 36 species of opisthokonts, including 26 animals (Meta36-Homo) using an irreversible Dollo binary substitution model. Singletons were excluded, and an ascertainment bias correction was applied for the absence gene families lost in all species. Numbers of nodes indicate posterior probabilities indistinguishable from 1.0. Convergence statistics are listed in Supplementary Table S1.

**Figure S7.**
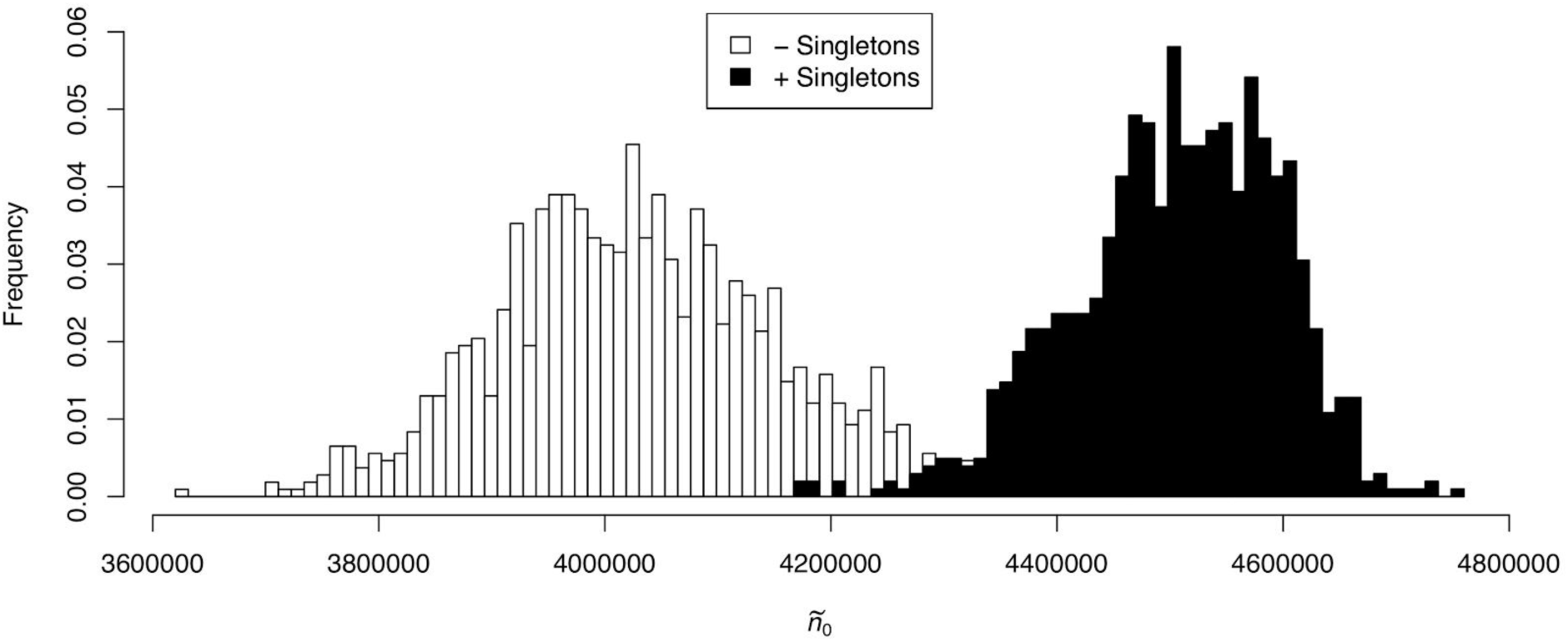
Posterior predictive distribution of the number of homologous gene families lost in all species (ñ_0_), inferred from Meta36-Ortho using a reversible binary substitution model, either including singletons (black) or excluding them (white).

**Figure S8.**
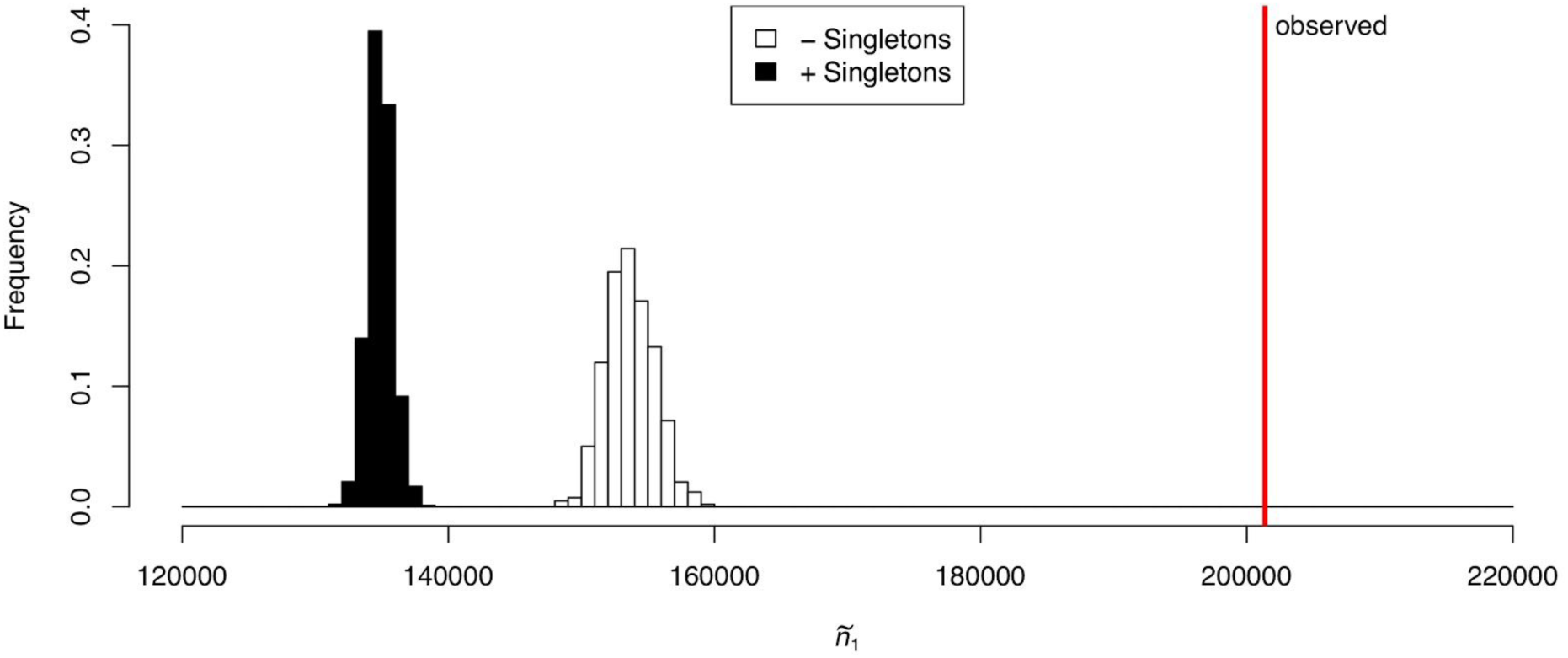
Posterior predictive distribution of the number of homologous gene families present in only a single species (ñ_1_), inferred from Meta36-Ortho using a reversible binary substitution model, either including singletons (black) or excluding them (white).

## Supplementary Table

**Table S1.** Convergence statistics computed using bpcomp and tracecomp

| **Taxa** | **Method** | **Singletons** | **Model** | **min effsize** | **max rel_diff** | **maxdiff** |
| --- | --- | --- | --- | --- | --- | --- |
| 23 | Ortho | No | Reversible | 762 | 0.076 | 0.009 |
| 23 | Homo | No | Reversible | 507 | 0.005 | 0.001 |
| 36 | Ortho | Yes | Reversible | 793 | 0.080 | 0.000 |
| 36 | Ortho | No | Reversible | 1306 | 0.099 | 0.031 |
| 36 | Homo | No | Reversible | 201 | 0.093 | 0.027 |
| 36 | Homo | Yes | Reversible | 399 | 0.072 | 0.009 |
| 36 | Homo | No | Dollo | 338 | 0.093 | 0.008 |

